# *Parageobacillus thermoglucosidasius* and the reductive glycine pathway: A journey of Almosts and Maybes

**DOI:** 10.1101/2025.04.15.648870

**Authors:** Miguel Paredes-Barrada, Stefano Donati, Richard van Kranenburg, Nico J. Claassens

## Abstract

We aimed to engineer the reductive glycine pathway (rGlyP) in *Parageobacillus thermoglucosidasius* for synthetic assimilation of formate and methanol via a mixed rational and evolutionary approach. We attempted to obtain glycine or serine auxotrophic strains in *P. thermoglucosidasius* as a starting point to impose a selective pressure and engineer formate assimilation via the rGlyP. While serine auxotrophy was partially achieved via gene deletion of *serA*, alternative glycine biosynthesis routes limited its stringency, and glycine auxotrophy could not be obtained due to unsuccessful genome modification. Even though formate supplementation improved the growth of the *serA* deficient strain, we confirmed that formate was not assimilated into biomass via ^13^C-labeling experiments. Thus, we hypothesized the increase in growth of the *serA* deficient mutant was likely due to extra energy provided by formate oxidation, rather than biomass incorporation. We also attempted heterologous overexpression of the C1 module from *Moorella thermoacetica*, but this did not yield any detectable effect in growth nor formate assimilation. These findings highlight the challenges in rGlyP implementation in non-model organisms, the need for better metabolic annotation in *P. thermoglucosidasius*, and provide valuable insights for future engineering of synthetic C1 metabolism in thermophiles via the rGlyP.

## Introduction

The use of microbial cell factories is a promising strategy to produce chemicals more sustainably. Several chemical products such as ethanol and lactate are already produced at a commercial scale by using this approach^1–5^. Sugar-based feedstocks are mostly used for the manufacturing of bio-based products, but there is an interest to move towards the use of alternative feedstocks to make bio-based manufacturing more sustainable and less dependent on agricultural land^6,7^. Alternative sustainable feedstocks should be generated from renewable resources and ideally not be dependent on agriculture, as this can potentially lead to competition with food production for production of bulk products, and a further reduction of the habitat for natural biodiversity.

Among alternative carbon feedstocks, one-carbon feedstocks such as formic acid, carbon monoxide and methanol are promising candidates, given that they can be produced from renewable resources. The use of thermophilic organisms for biobased manufacturing can reduce the amount of energy needed to cool the bioreactors, as well as the risk of contamination by mesophilic organisms^8^. Thus, enabling synthetic assimilation of methanol or formate in a thermophilic and versatile microorganism represents an important step in utilizing C1 compounds for sustainable biomanufacturing. *Parageobacillus thermoglucosidasius* is a suitable thermophilic bacterial chassis for bioproduction from C1 feedstocks, given its expanding genetic toolbox, metabolic versatility and robustness in growth. In fact, in our previous study we demonstrated its potential as a bacterial chassis by engineering partial methylotrophy via the RuMP cycle in this thermophile^9,10^.

Apart from the RuMP cycle, other C1 assimilating pathways exist in nature or have been engineered by humans^11,12^. Therefore, alternative pathways could be explored to enable synthetic methanol or formate assimilation in *P. thermoglucosidasius*. The reductive glycine pathway (rGlyP) stands out among other aerobic C1 assimilating pathways because of its energetic and kinetic efficiency, and has demonstrated success in enabling (partial) formate or methanol assimilation in engineered strains of *Cupriavidus necator, Escherichia coli, Pseudomonas putida* and *Sacharomyces cerevisiae*^11,13–17^. In this study, we attempted to implement the rGlyP in *P. thermoglucosidasius* to enable synthetic utilization of formate or methanol via this pathway.

Initially designed as a synthetic C1 assimilation pathway, an anaerobic variant of rGlyP was later found to occur naturally in *Desulfovibrio desulfurican*s^18^. When comparing the rGlyP to other aerobic formate assimilation pathways, such as the naturally occurring serine cycle or the Calvin cycle, the rGlyP demonstrates superior efficiency in terms of less ATP requirements^19^. Moreover, based on current pathway kinetics data and thermodynamic driving forces, the rGlyP is recognized as one of the most effective pathways for formate assimilation^11,19^. In contrast, when evaluating methylotrophic pathways, the data does not clearly establish the rGlyP as superior. For example, using the RuMP cycle as a benchmark, the rGlyP demands more ATP per unit of pyruvate or biomass produced, being less efficient. Regarding pathway kinetics, which may have a greater impact on growth and production rates than ATP requirements, current data suggest that the rGlyP pathway kinetics could be comparable to those of the RuMP cycle^6,19^.

Another interesting aspect of the rGlyP is that it requires the co-assimilation of CO_2_ through the reversed (reducing) glycine cleavage system, with methanol or formate as co-substrates, and this presents both advantages and disadvantages. On one hand, a CO_2_ rich atmosphere is required, which could pose a challenge for bioprocessing, as it would require additional infrastructure compared to using methanol or formate as the sole substrate, such as extra gas tanks and pipelines. However, when methanol is the sole substrate, its high reduction state presents a challenge for targeting oxidized products. To maintain redox balance, some NADH derived from methanol must be dissipated through respiration or the production of reduced by-products, leading to a decrease in product yield. Because CO_2_ reduction acts as an electron sink, a pathway that co-utilizes methanol and CO_2_ can therefore increase the yield of methanol on oxidized products like e.g., lactate^11^. Furthermore, CO_2_ can be a cheaper substrate than methanol or formate when nearby industrial pure CO_2_ waste-streams are available, having in some contexts even a negative price^20,21^. Therefore having an input of carbon feedstock coming from mixed CO_2_ and either formate or methanol, could be interesting from an economic point of view.

Notably, when formate is used as the only carbon source, ~84% of the formate need to be oxidized to produce enough reducing power to drive the assimilation of ~16% of the formate^11^. CO_2_ is produced as a result of formate oxidation (Figure 1), and depending on the bioreactor characteristics, this could alleviate or totally avoid the need to feed extra CO_2_ to the bioreactors.

**Figure 1:**
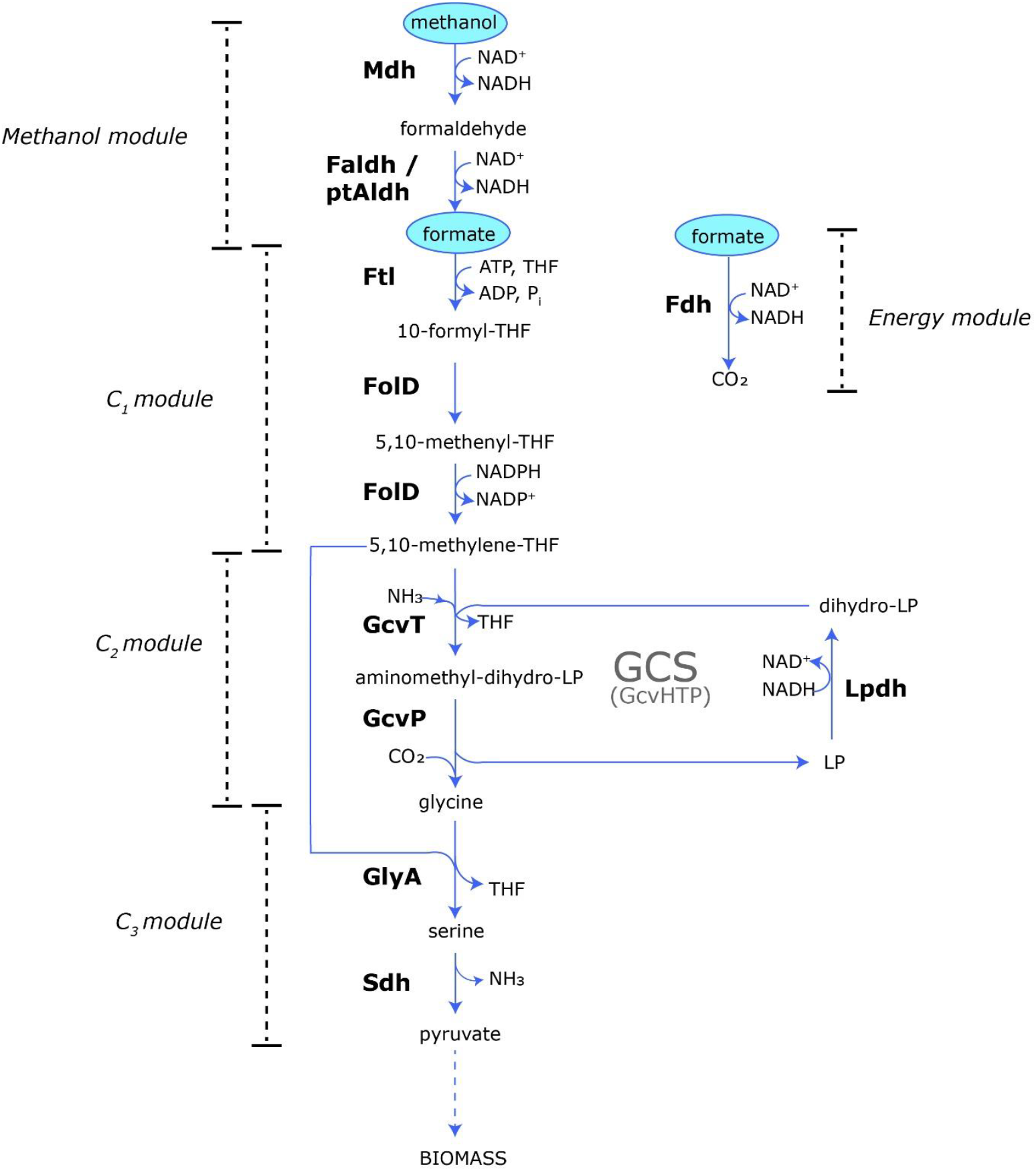
The rGlyP. Possible substrates of the pathway, methanol or formate, are in blue ovals. Dashed lines represent multiple reactions. GCS: Glycine cleavage system. Mdh: methanol dehydrogenase. Faldh: formaldehyde dehydrogenase. Fdh: formate dehydrogenase. PtAldh: putative formaldehyde dehydrogenase native to *P. thermoglucosidasius*. Ftl: formate-tetrahydrofolate (THF) ligase. FolD: bifunctional 5,10-methenyl-THF cyclohydrolase, 5,10-methylene-THF dehydrogenase. GcvT: Protein T of the glycine cleavage system complex. GcvP: Protein P of the glycine cleavage system complex. GcvHTP: Glycine cleavage complex. LpdH: lipoamide dehydrogenase. GlyA: serine hydroxymethyltransferase. Sdh: serine dehydratase.

Moreover, the rGlyP is a linear pathway that does not overlap with central carbon metabolism, such as glycolysis or the pentose phosphate pathway. This makes it potentially easier to engineer in a microorganism, as central metabolism is often intricately regulated^22^. However, the successful engineering of pathways that overlap with central carbon metabolism demonstrates that this challenge can be overcome, e.g., the RuMP cycle^10,23–25^. All the aforementioned features make the rGlyP an attractive pathway for enabling C1 feedstock assimilation in the emerging thermophilic model organism *P. thermoglucosidasius*.

The rGlyP (Figure 1) uses the cofactor tetrahydrofolate (THF) to condense formate into formyl-THF, via the formate THF ligase (Ftl). Formyl-THF then gets dehydrated and reduced to 5,10-methylene-THF, in a series of reactions that can be catalyzed by a bifunctional enzyme that encompasses both cyclohydrolase and dehydrogenase activities, named FolD. 5,10-methylene-THF is incorporated into the glycine cleavage system (GCS), which can be forced to work in the reductive direction in a CO_2_ rich environment, yielding a molecule of glycine. In the canonical rGlyP variant, the enzyme serine hydroxymethyltransferase (GlyA) converts glycine into serine via condensation with another C1 moiety, coming from 5,10-methylene-THF. Serine enters central carbon metabolism through deamination to pyruvate, via the serine dehydratase (Sth). In this way, the rGlyP assimilates formate into glycine, which then gets assimilated into the metabolism via serine and pyruvate (Figure 1).

The rGlyP also possesses an energy module, in which formate is oxidized to CO_2_ and NADH is produced, in a reaction catalyzed by the formate dehydrogenase (Fdh). Additionally, a methanol module can be added to enable methanol assimilation via the rGlyP^14,16^. This includes the oxidation of methanol to formaldehyde, via a methanol dehydrogenase (Mdh) and the subsequent oxidation of formaldehyde to formate, via a formaldehyde dehydrogenase (Faldh)^11,14^. Formaldehyde dehydrogenases, like other dehydrogenases, are dependent on cofactors such as glutathione (mainly present in Gram-negative bacteria), bacillithiol (BSH; mostly present in Gram-positive bacteria), NAD^+^ or others, to catalyze redox reactions, typically taking part in native detoxification pathways^26^. In bacterial species from the *Bacillaceae* family, such as *B. subtilis*, different BSH-dependent and independent formaldehyde dehydrogenases are present, which typically are involved in formaldehyde detoxification pathways among other enzymes^26–31^.In evolved strains of *P. thermoglucosidasius* partially growing on methanol via the RuMP, we observed that the enzyme PtAldh, an uncharacterized aldehyde dehydrogenase native to *P. thermoglucosidasius*, was drastically underproduced in the evolved partially methylotrophic PTM1 strain. Thus, we hypothesized that PtAldh could be a formaldehyde dehydrogenase, which takes part in a formaldehyde detoxification pathway^10^. If this assumption is correct, this enzyme would be able to catalyze the oxidation of formaldehyde and possibly be used in the rGlyP methanol module to convert formaldehyde into formate (Figure 1).

Examples of engineering the rGlyP have focused on a modular approach, by which different modules of the pathway are engineered in a sequential way^11,14,16,17^. The rGlyP pathway is organized into distinct modules: the energy module, which includes the reaction catalyzed by Fdh and generates energy for the cell; the C1 module, encompassing the reactions that convert formate to 5,10-methylene-THF; the C2 module, comprising the reactions that transform 5,10-methylene-THF into glycine; the C3 module, which involves the conversion of glycine to pyruvate; and the methanol module, responsible for the oxidation of methanol to formaldehyde and subsequently to formate (Figure 1). Typically, the rGlyP is first engineered to partially assimilate formate via some of the modules, then optimized by adaptive laboratory evolution (ALE) to assimilate formate as a sole carbon source and the last module to be introduced is the methanol module, which converts methanol to formate ^11,14,16,17^.

Implementing synthetic C1 assimilation is a challenging process, which implies a holistic rewiring of central carbon and energy metabolism, as well as a readjustment of cellular homeostasis. Because of such complexity, typically efforts to engineer the rGlyP relied on a mixed approach, including rational engineering and ALE. These mixed approaches are often based on rational engineering of formate dependent strains as a starting point for ALE, and it is combined with the sequential implementation of the different modules ^11,14,16,17^. Formate dependence can be achieved by one or more gene knockouts that make the cells unable to grow solely on a certain auxiliary carbon substrate, unless they co-assimilate formate. Such deletion strains are then normally complemented with heterologous enzymes that enable formate assimilation, such as a Fdh, Ftl, FolD and others. Given that the GCS naturally operates in the oxidative direction to provide cell metabolism with C1 moieties from glycine, forcing the GCS to work in the reductive direction in a CO_2_-rich environment, is also required to successfully engineer the rGlyP ^11,14,16,17^. After formate dependency and heterologous enzymes are introduced, ALE is typically performed to enhance functional operation of the rGlyP under CO_2_ rich conditions.

The modular engineering approach of the rGlyP enables various selection strategies with differing levels of stringency, requiring a greater or lesser percentage of biomass to be derived from formate assimilation. This, in turn, imposes lower or higher selection pressures on the engineered strains. Low selection pressure strategies include the C1 auxotroph (Δ*gcvTHP*, Δ*glyA*; ~3% biomass originating from formate, when supplemented with glycine) and the C1 + β-carbon serine auxotroph (Δ*gcvTHP*, Δ*serA*; ~4% biomass originating from formate, when supplemented with glycine)^11^. These strategies were primarily used as proof of concept for the C1 module of the rGlyP during its early development in *E. coli* ^11,32,33^. Higher selection pressure strategies, which require more biomass coming from formate assimilation, like the glycine or serine auxotrophy, have been more commonly applied to achieve synthetic formatotrophy or methylotrophy via the rGlyP in engineered microorganisms (Figure 2)^11^.

**Figure 2:**
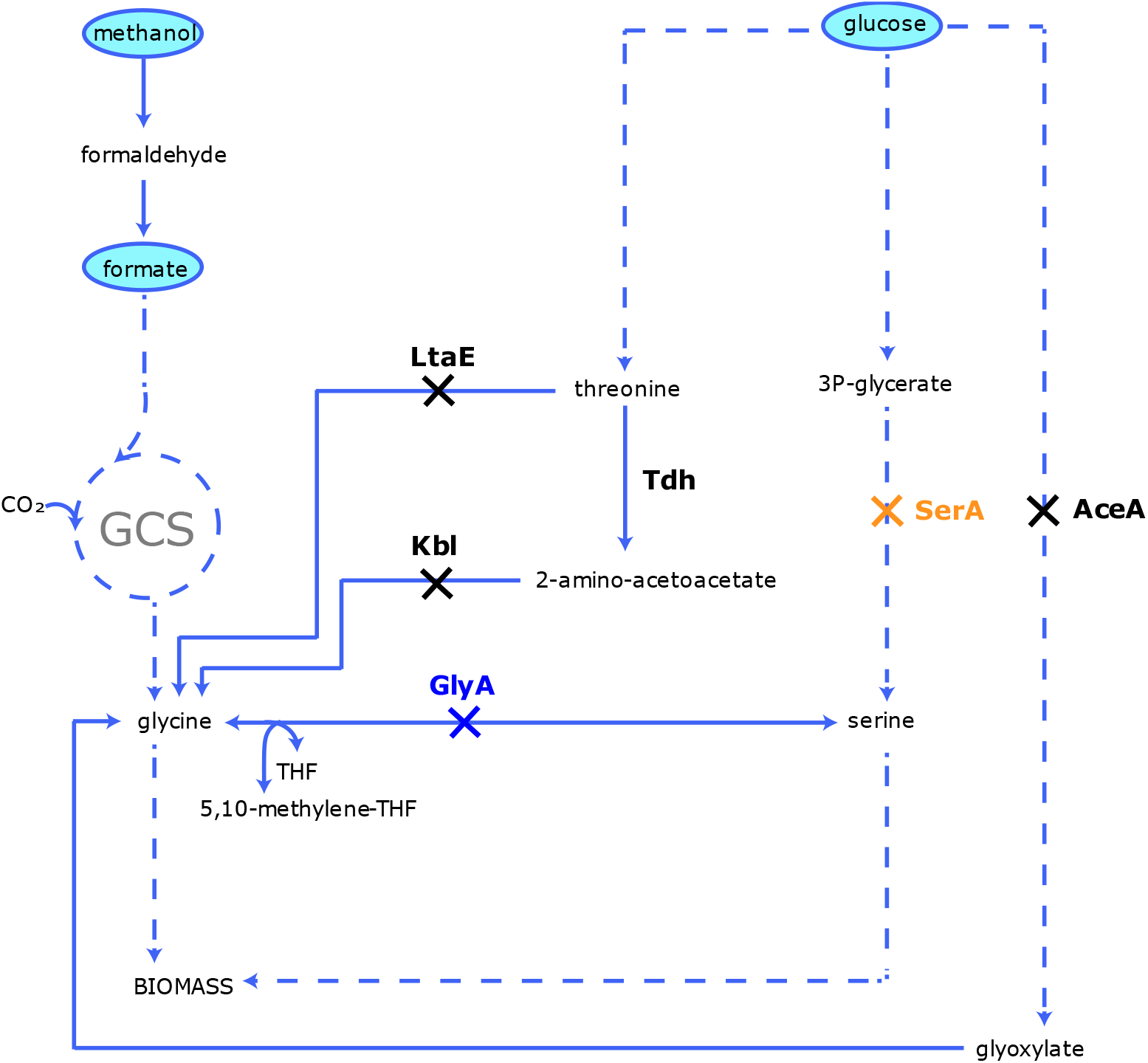
Most used selection strategies employed to engineer the C1 module of the rGlyP in different microorganisms, the glycine auxotrophic and serine auxotrophic strains. The glycine auxotrophic strains, involves the disruption of glycine biosynthesis from threonine (Δ*ltaE, Δkbl*), serine (Δ*glyA*) and glyoxylate (Δ*aceA*), while the serine auxotrophic strains involve the disruption of glycine biosynthesis from threonine (Δ*ltaE*, Δ*kbl*) and glyoxylate (Δ*aceA*), as well as the disruption of serine biosynthesis from 3PG (Δ*serA*). Dashed lines represent multiple reactions. GCS: glycine cleavage system. GlyA: serine hydroxymethyltransferase. SerA: 3PG dehydrogenase. AceA: isocitrate lyase. Tdh: threonine dehydrogenase. Kbl: glycine acetyltransferase. LtaE: l-threonine aldolase

The selection through glycine auxotrophic strains is based on the disruption of the native pathways leading to glycine biosynthesis, from serine (Δ*glyA*), from threonine (Δ*ltaE*, Δ*kbl*) and/or from glyoxylate via transaminases (Δ*aceA*), and requires that ~8% biomass is assimilated through the C1 module of the rGlyP^11^. This selection strategy has been applied to engineer the rGlyP in *C. necator* and *E. coli*, and required overexpression of the C1 and C2 module enzymes^13,14,34^. Following this, reconstitution of GlyA was necessary to achieve full formatotrophy, which was further improved through an additional round of ALE to optimize the entire pathway.

The selection through serine auxotrophic strains involves the disruption of glycine biosynthesis from threonine (*ΔltaE* and Δ*kbl*) and serine biosynthesis from 3-phosphoglycerate (Δ*serA*), and requires that 11% biomass is assimilated from formate^11^. The serine auxotrophic selection scheme has the advantage over the other selection schemes that none of the disrupted enzymes are within the rGlyP, which avoids the need for reconstituting the deleted genes after ALE to achieve full formatotrophy. Engineering the rGlyP in *E. coli* via the serine auxotrophic selection scheme successfully yielded a full formatotrophic strain, which required the overexpression of the C1 module and GCS enzymes, but not of GlyA^34^. This strategy also enabled synthetic formatotrophy in *P. putida*^16^.

As previously mentioned, enabling synthetic methylotrophy via the rGlyP is possible^14,16^. The methanol module encompasses the conversion of methanol to formate, catalyzed by methanol dehydrogenase (Mdh) and formaldehyde dehydrogenase (Faldh). A key study demonstrated that overexpression of a heterologous methanol dehydrogenase is essential for engineering the methanol module within the rGlyP pathway, while the native formaldehyde dehydrogenase in *E. coli* (encoded by *frmAB*) proved sufficient for efficient formaldehyde oxidation^14^. Various methanol dehydrogenases have been evaluated for this purpose; however, the only enzyme capable of supporting methylotrophic growth was the methanol dehydrogenase from *Geobacillus stearothermophilus* (BsMdh)^14^. This achievement led to a methylotrophic *E. coli* strain capable of growing on methanol with a doubling time of ~55 hours, highlighting the need for further optimization.

In this study, we aimed to enable formate and methanol assimilation via the rGlyP in *P. thermoglucosidasius* using a mixed rational approach based on the selection of glycine and serine auxotrophs. By using the ThermoCas9 counterselection system, we successfully deleted the *serA* gene but were unable to obtain the Δ*glyA* mutant^35^. Although the selection pressure in the Δ*serA* strain was not entirely stringent, promising results were observed in cultures supplemented with formate and CO, leading us to hypothesize that the C1 module of the rGlyP was active. However, steady-state ^13^C-labeling experiments with labeled formate revealed that formate was not being assimilated into glycine, invalidating this hypothesis. Despite this, we believe these results offer valuable insights for engineering the rGlyP pathway in *P. thermoglucosidasius*.

## Results and discussion

### Implementing the selection schemes of glycine and serine auxotrophic strains in *P. thermoglucosidasius*

Creating glycine and serine auxotrophic strains has proven to be an effective strategy for selecting a functional rGlyP C1 module and enabling formate assimilation in various microorganisms^11,13,14,16,17,32,34^. These selection schemes typically involve disrupting native pathways responsible for glycine biosynthesis. For glycine auxotrophs, this requires inactivating pathways that produce glycine from serine (*ΔglyA*), threonine (*ΔltaE, Δkbl*), and glyoxylate (*ΔaceA*). For serine auxotrophs, it is necessary to disrupt pathways that natively produce serine (*ΔserA*) and convert threonine and glyoxylate to glycine (Figure 2).

We aimed to develop modified strains of *P. thermoglucosidasius* using these two parallel strategies, the glycine and serine auxotrophy selection schemes. However, due to the poor annotation of the *P. thermoglucosidasius* genome, the pathways converting threonine to glycine are not (yet) described. Initially, we hypothesized that these pathways might be absent, thus we focused on deleting specific genes producing glycine from serine (*glyA*), or the biosynthesis of serine itself (*serA*). Once one of these pathways was disrupted, we also aimed to obtain a double mutant with an *aceA* deletion to block glyoxylate production and subsequent conversion to glycine via amino transferases.

To achieve this, we constructed editing plasmids containing 1 kb homology arms flanking the target genes *glyA* (BCV53_03760), *serA* (BCV53_RS15865) and *aceA* (BCV53_RS06880), and the ThermoCas9 counterselection tool to obtain the single mutants Δ*glyA*, Δ*serA* and double mutants Δ*glyA* Δ*aceA* and Δ*serA* Δ*aceA* strains of *P. thermoglucosidasius* (Table 2)^35^. Despite several attempts, we only achieved the *serA* gene deletion and were unable to obtain the Δ*glyA* mutant. Consequently, the glycine auxotrophic selection scheme was discarded for this project. Moreover, we also failed to disrupt glyoxylate biosynthesis in the Δ*serA* strain, thus we could not achieve the double mutant strain Δ*serA* Δ*aceA*.

Having obtained the *P. thermoglucosidasius* Δ*serA* strain, we aimed to investigate how stringent the disruption of the pathway leading to serine biosynthesis, and ultimately to glycine via GlyA, was to select for the C1 module of the rGlyP. Optimally, to ensure stringent selection of the C1 module via the serine auxotrophy, total abolishment of growth on glucose is desired. Therefore, we set up growth experiments in falcon tubes, in which we tested if the Δ*serA* mutant was able to grow on glucose as a sole carbon source in minimal media. We also included either glycine or serine supplementation as positive controls to cultures of *P. thermoglucosidasius* Δ*serA* and WT strains with glucose as the sole carbon source (Table 1). We included glycine supplementation because we hypothesized that addition of glycine could also rescue the growth of the Δ*serA* strain through its conversion to serine via GlyA.

**Table 1:**
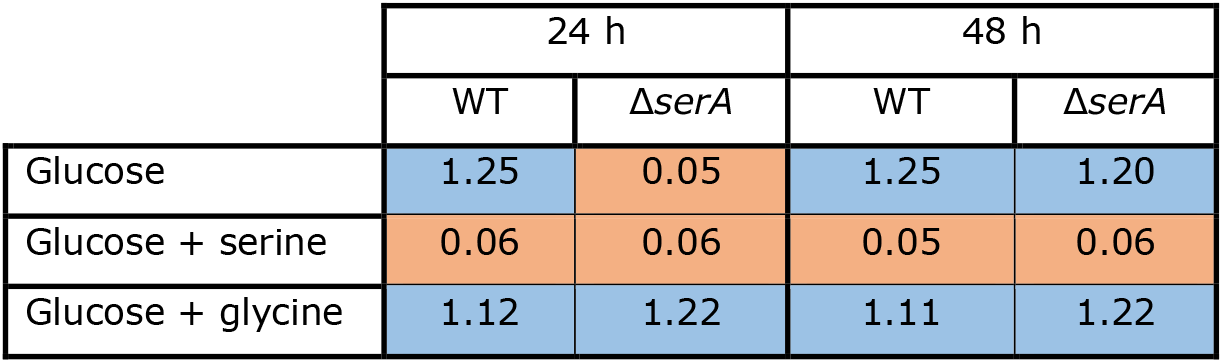
Average OD_600_ of triplicate cultures of *P. thermoglucosidasius* WT or Δ*serA* after 24 or 48 h, when glucose (2 g/L) was used as the sole carbon source, or when supplemented with either glycine (150 mg/L) or serine (210 mg/L). Blue cells = growth. Orange cells = no growth.

If the disruption of serine biosynthesis completely blocked glycine production, and *P. thermoglucosidasius* lacked both alternative pathways to synthesize glycine from threonine and efficient pathways to derive it from glyoxylate, as we hypothesized, then no growth would be expected in the *ΔserA* mutant when cultivated on glucose.

While growth of the *P. thermoglucosidasius* Δ*serA* strain was abolished during the first 24 h, and only rescued by the addition of glycine, the mutant strain could grow up to an OD_600_ comparable to that of the WT strain after 48 h. Notably, neither the *P. thermoglucosidasius* WT or Δ*serA* strains were able to grow in the presence of serine, indicating that high concentrations of serine might be toxic to the cells, which is actually a known phenomenon^36,37^.

These results indicated that alternative routes leading to glycine biosynthesis, other than from serine, were operating in *P. thermoglucosidasius*. Known metabolic routes leading to glycine biosynthesis other than from serine, take either threonine or glyoxylate as substrates^11^. In other microorganisms, such as the *P. thermoglucosidasius* close relative *B. subtilis*, two pathways converting threonine to glycine can be found. One that uses the L-threonine aldolase (LtaE) to convert threonine to glycine in a single enzymatic step, and another one that is catalyzed by two enzymes, threonine dehydrogenase (Tdh) and glycine acetyltransferase (Kbl; Figure 3)^38^. Despite our efforts to find the homologous enzymes to LtaE, Tdh and Kbl in *P. thermoglucosidasius* via BLAST searches, we could not find the homologous enzymes that would catalyze the conversion of threonine to glycine.

**Figure 3:**
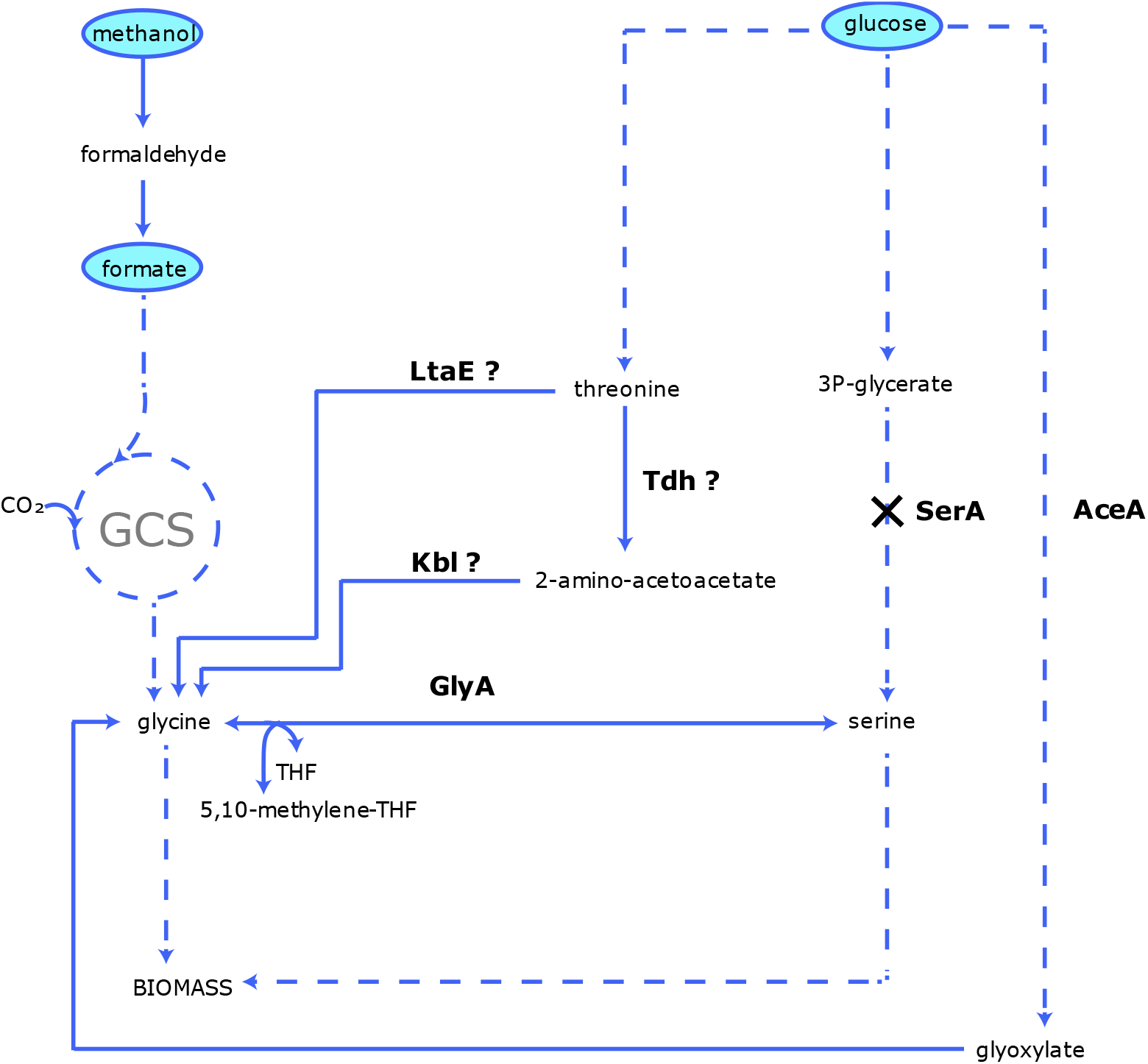
Metabolic landscape of the *P. thermoglucosidasius* Δ*serA* strain. GCS: glycine cleavage system. Dashed lines represent multiple reactions. GlyA: serine hydroxymethyltransferase. SerA: 3PG dehydrogenase. AceA: isocitrate lyase. Tdh: threonine dehydrogenase. Kbl: glycine acetyltransferase. LtaE: L-threonine aldolase

In relation to the conversion of glyoxylate to glycine, it is a reaction that can occur via different amino transferases. Typically, microorganisms possess many of these enzymes (*P. thermoglucosidasius* has more than 10), and therefore deleting all of them can become difficult and will probably render a phenotype with reduced fitness^11,14^. Thus, a better way to impede glycine biosynthesis from glyoxylate in *P. thermoglucosidasius* is to block glyoxylate biosynthesis by deleting the gene encoding AceA^11,14^. However, despite several attempts, we were unable to obtain a Δ*aceA* mutant strain of *P. thermoglucosidasius*.

Our failure to construct the Δ*serA* Δ*aceA* strain, combined with the limited genomic and metabolic information available for *P. thermoglucosidasius*, hindered our understanding of why the Δ*serA* strain was still able to grow on glucose.

### Heterologous overexpression of the C1 module from *Moorella thermoacetica* had no effect on growth, but formate addition improved growth of the Δ*serA* mutant

We were unable to generate a fully stringent *P. thermoglucosidasius* strain for selecting the C1 module. However, we hypothesized that the growth delay of the Δ*serA* mutant when glucose is used as the sole carbon source (Table 1), could still create sufficient growth disparity to confer a selective advantage for formate assimilation via the C1 module of the rGlyP.

We hypothesized that formate conversion into glycine could probably rescue the impaired growth phenotype of the Δ*serA* mutant, in the same way as glycine supplementation does. Furthermore, overexpression of the C1 module in the *P. thermoglucosidasius* Δ*serA* strain, could potentially further increase the potential of formate assimilation into glycine, under a CO_2_ rich atmosphere. For that purpose, we chose to heterologously overexpress the Ftl and FolD enzymes from the thermophilic acetogen *M. thermoacetica*^*39–41*^. We built the plasmid pJS15A-Ftl-FolD-MT and transformed it in *P. thermoglucosidasius* WT and Δ*serA* strains, yielding the *P. thermoglucosidasius* WT-C1MT and Δ*serA*-C1MT strains (Figure 4A; Table 2). We also transformed the pJS15A-EV (empty vector) into *P. thermoglucosidasius* WT and Δ*serA*, yielding the strains *P. thermoglucosidasius* WT-EV and Δ*serA*-EV, which served as negative controls (Table 2).

**Table 2:**
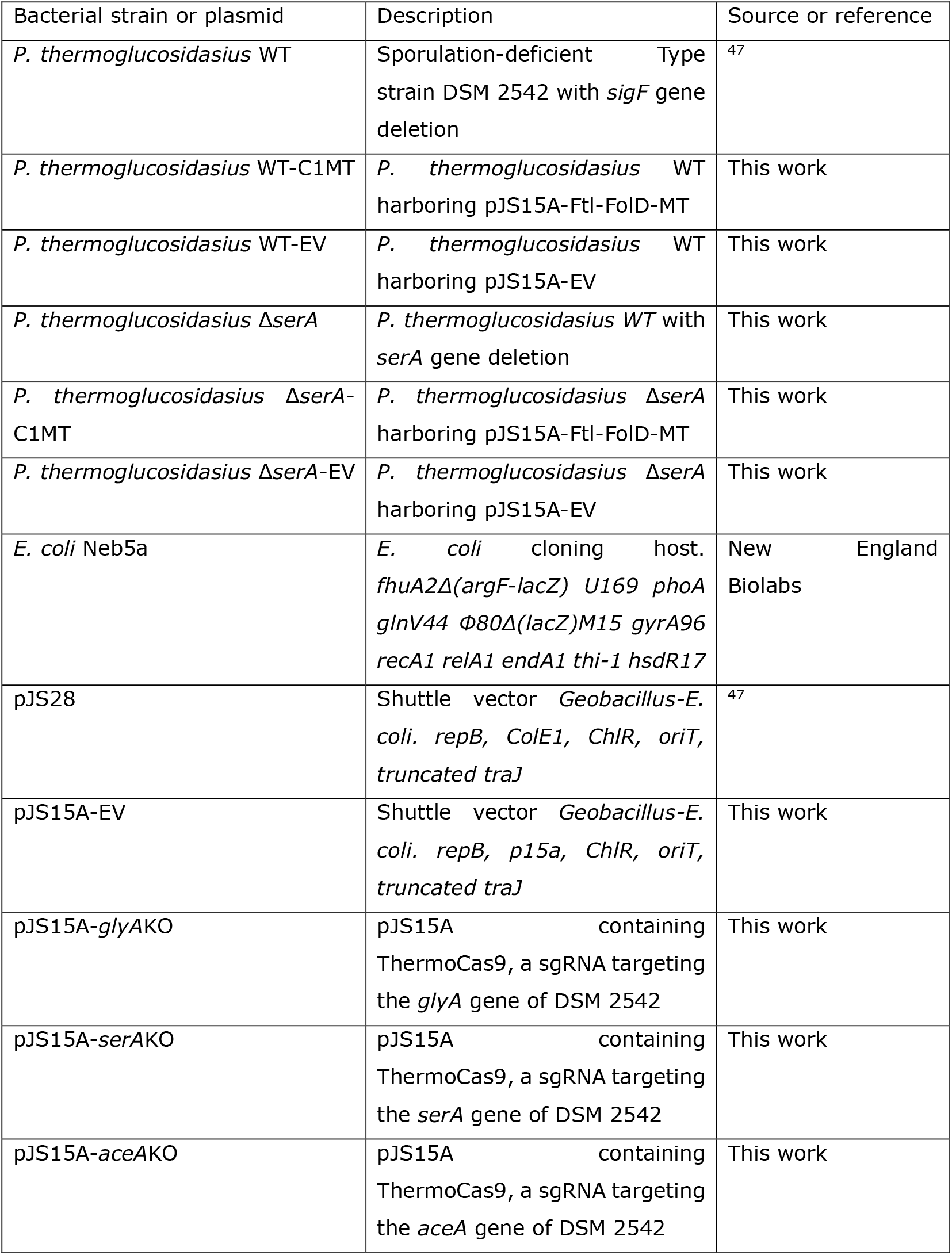
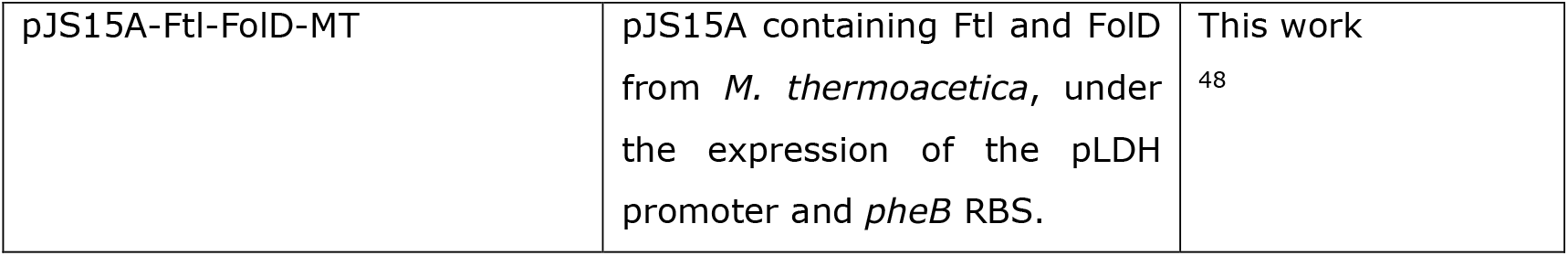
List of bacterial strains and plasmids used in this study, together with a brief description and the source.

**Figure 4:**
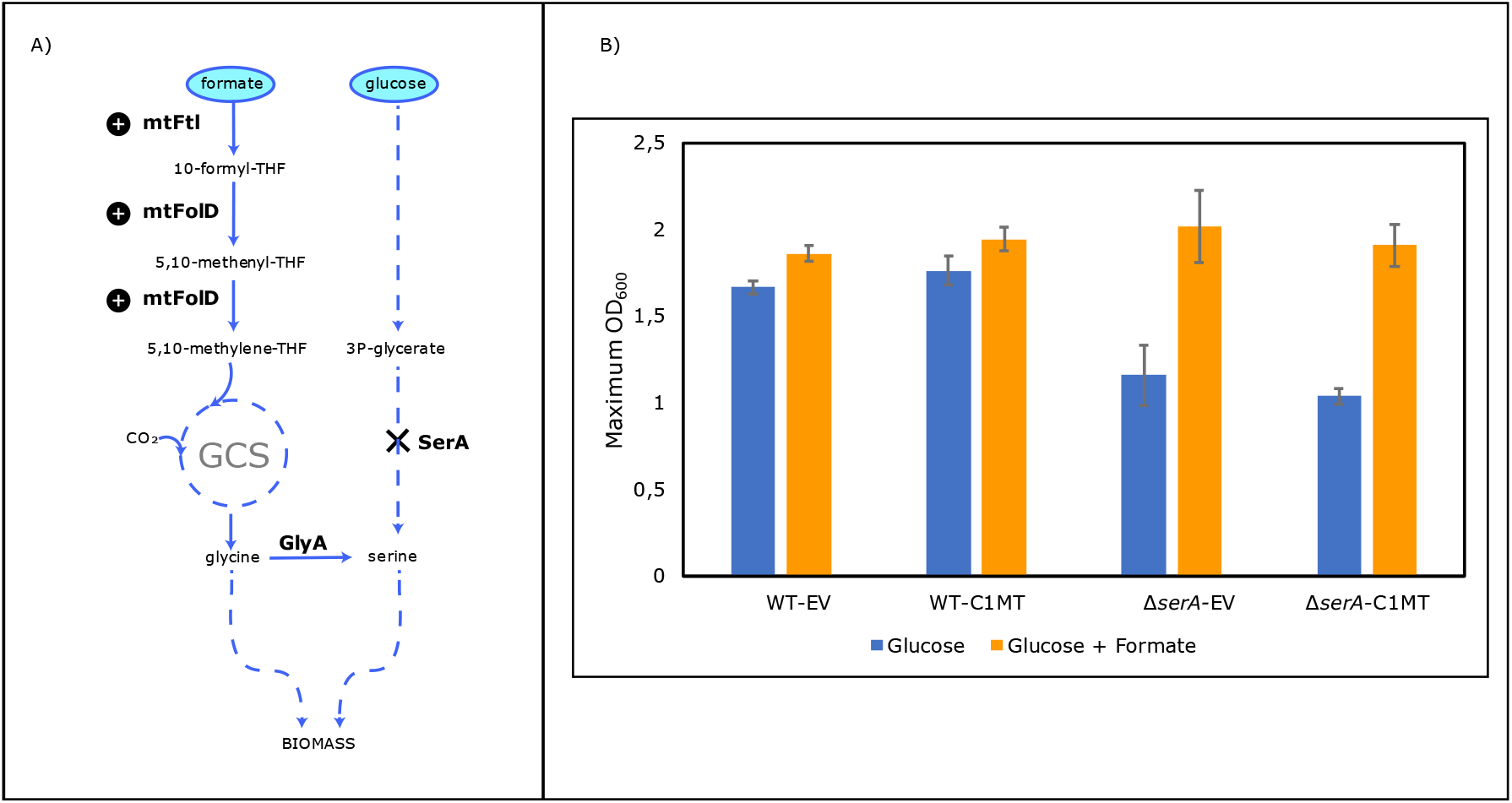
A) C1 module overexpression in the *P. thermoglucosidasius* Δ*serA*-C1MT strain. (+) symbols represent heterologously overexpressed enzymes from *M. thermoacetica*. Dashed lines represent multiple reactions MtFtl= formate-tetrahydrofolate ligase from *M. thermoacetica*. MtFolD= bifunctional 5,10-methenyl-THF dehydratase, 5,10-methylene-THF dehydrogenase, from *M. thermoacetica*. SerA= 3PG dehydrogenase. GlyA= serine hydroxymethyltransferase. B) Maximum OD_600_ averages of cultures of *P. thermoglucosidasius* WT-EV, WT-C1MT, *ΔserA*-EV and *ΔserA*-C1MT growing in minimal medium with CO_2_(10%) and either glucose (5 g/L) or a mixture of glucose (5 g/L) and formate (1.35 g/L). Error bars represent standard deviation for five replicate cultures.

To determine whether formate was assimilated via the rGlyP and influenced the growth of *P. thermoglucosidasius* WT-EV, WT-C1MT, Δ*serA*-EV or Δ*serA*-C1MT, we performed growth experiments in minimal media containing either glucose (5 g/L) as the sole carbon source or a combination of glucose (5 g/L) and formate (1.35 g/L), as well as CO_2_ (10%; Figure 4B).

The Δ*serA* EV strain exhibited growth impairments when cultivated solely on glucose, reaching a 44% lower maximum optical density than the WT-EV strain. This was also observed for Δ*serA*-C1MT, which reached a 70% lower maximum OD_600_ than the WT-C1MT strain. As observed previously, the *ΔserA*-EV and *ΔserA*-C1MT strains exhibited an extended lag phase of ~24 hours, which was not alleviated by the addition of formate to the medium (Figures S1-S4). However, the addition of formate to the medium increased the maximum OD_600_ of the Δ*serA* and Δ*serA*-C1MT strains, achieving a comparable growth to the WT-EV and WT-C1MT strains and improving their maximum OD_600_ by 74% (Δ*serA*) and 84% (Δ*serA*-C1MT) compared to their growth on glucose alone. This suggests that formate has a positive effect on the impaired growth of the serine autoxtrophic strains. The presence of formate did not have a big effect on maximum OD_600_ of the WT-EV or WT-C1MT strains (Figures 4B, S1-S4).

However, the overexpression of the C1 module did not significantly enhance growth in either the WT or Δ*serA* mutant strains, suggesting that formate was not assimilated via the heterologously expressed enzymes from *M. thermoacetica* (Figure 4B, S1-S). Instead, we hypothesized that the restored growth of the Δ*serA* strain in the presence of formate could be attributed to its assimilation into glycine through the native *P. thermoglucosidasius* enzymes Ftl and FolD. To test this hypothesis, we conducted ^13^C steady-state experiments to validate formate assimilation via this native pathway.

### ^13^C labelling experiments showed that the C1 module of the rGlyP was not active in *P. thermoglucosidasius*

To validate formate assimilation into biomass, we performed labelling experiments with ^13^C-labelled formate. The WT-C1MT, Δ*serA*-EV and Δ*serA*-C1MT were cultivated in minimal TMM medium with ^12^C-glucose (5 g/L), ^12^C-CO_2_ (10%), and ^13^C-labelled formate (1.38 g/L). Cell pellets were analyzed to investigate the incorporation of the 13C-labelled formate in the proteogenic amino acids glycine and serine. We hypothesized that if formate was co-assimilated with glucose and CO_2_ via the rGlyP enzymes, one of the carbon atoms of glycine should be labelled, as well as two of the carbons of serine (Figure 5A). Taking into account that the selection posed by the *serA* deletion was not fully stringent and some serine and glycine could still be synthesized from glucose via unknown pathways, it was difficult to know which would be the expected percentage of labelled amino acid molecules. However, in the case of formate assimilation via the rGlyP, surely there should be a significant percentage of glycine and serine labelled molecules.

**Figure 5:**
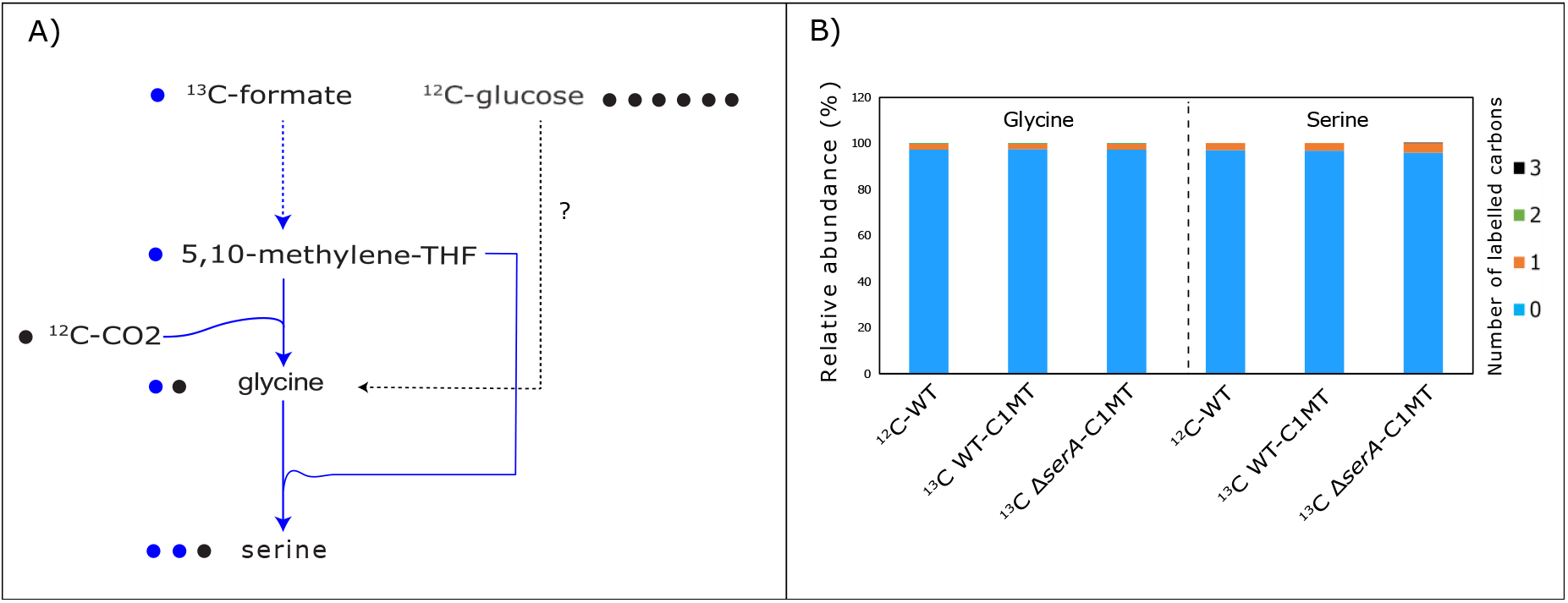
A) Metabolic scheme showing the expected labelling pattern of glycine and serine assuming co-assimilation of ^13^C-labelled formate, ^12^C-CO_2_ and ^12^C-glucose. B) Relative abundance of ^13^C-labelled carbons present in the proteogenic amino acids glycine and serine, in cells of the WT-C1MT and Δ*serA*-C1MT strains cultivated in the presence of a mixture of natural ^12^C glucose, ^12^C CO_2_ and ^13^C-labelled formate. The WT strain cultivated on natural ^12^C ribose was used as control to account for naturally occurring ^13^C atoms.

The results of the analysis revealed that neither glycine nor serine had any labelled carbon in any of the analyzed strains (Figure 5B). This was a surprising result to us given the differences in maximum OD_600_ observed in the growth experiments with formate addition (Figure 4B), but it was clear evidence that formate was not being assimilated via the rGlyP in any of the tested strains.

This result also raises the question of why the Δ*serA*-C1MT strain exhibited an 84% increase in maximum OD_600_ upon the addition of formate if this formate was not being assimilated into biomass (Figure 4B, 5B). One possible explanation is that the formate was oxidized to CO_2_ and NADH via the native Fdh of *P. thermoglucosidasius*, providing additional energy to the cells and partially alleviating the impaired growth phenotype of the Δ*serA*-EV and Δ*serA*-C1MT strains. This hypothesis seems counterintuitive, as these strains should be able to generate sufficient energy from glucose metabolism via glycolysis and the TCA cycle, making the extra NADH from formate seemingly unnecessary. However, it is possible that the pathway used by the *ΔserA* strains to synthesize serine has a higher energy demand, and the oxidation of formate may supply the necessary energy to support normal growth levels.

## Conclusion

The reductive glycine pathway (rGlyP) is a promising metabolic route for enabling synthetic formate/methanol assimilation in microorganisms^11^. In this study, we aimed to engineer the rGlyP in *P. thermoglucosidasius* using a mixed rational approach. This included generating glycine or serine auxotrophic strains and optimizing the pathway through ALE^11^. Although we were not successful in generating Δ*glyA* mutant strains obtained the Δ*serA* strain. However, we observed that this modification did not impose sufficient selection pressure to strictly select for formate assimilation, as the strain could still grow on glucose, although after an extended lag phase. This suggested the presence of alternative serine biosynthetic routes, likely involving glycine derived from either threonine or glyoxylate^38^. While glyoxylate derived glycine production is known in *P. thermoglucosidasius*, leaving this pathway intact has not hindered successful rGlyP engineering in other species^13,38^. Therefore, we propose that an uncharacterized glycine production pathway from threonine in *P. thermoglucosidasius* may contribute significantly to glycine biosynthesis, alongside the serine biosynthesis pathway that we disrupted. Additionally, glyoxylate transamination to glycine could also play a role in this process. The incomplete annotation of the *P. thermoglucosidasius* genome highlights the critical role of accurate metabolic knowledge in achieving successful metabolic engineering goals.

To investigate the hypothesized threonine-to-glycine pathway, growth experiments with the *ΔserA* strain could be conducted using glucose as the sole carbon source, supplemented with threonine. If threonine supplementation rescues growth similarly to glycine, it would provide preliminary evidence for this unknown pathway (Table 1). Further investigation could involve ^13^C-labeling experiments with labelled threonine or enzymatic assays. To investigate if the route from glyoxylate to glycine via AceA is efficient enough to rescue the growth of the *ΔserA* strain, a similar test with glucose and glyoxylate could also be conducted.

We also heterologously expressed the C1 module enzymes, Ftl and FolD, from *M. thermoacetica*. However, this did not result in detectable formate assimilation, potentially due to issues with protein expression, folding, or activity in *P. thermoglucosidasius*^*39*,*42*^. To determine whether the heterologous proteins of the C1 module are correctly expressed, an SDS-PAGE analysis of crude extracts from the overexpression strains could be performed. If the heterologous Ftl and FolD were successfully expressed, the native *P. thermoglucosidasius* Ftl and FolD enzymes could be deleted, allowing for formate complementation assays to assess whether the heterologous enzymes are correctly folded and functionally active. However, this approach would require a fully stringent selection strain in which all pathways leading to serine synthesis are disrupted.

Despite this C1 heterologous module was not expressed or inactive, the addition of formate to Δ*serA* cultures significantly increased maximum OD_600_, initially suggesting formate assimilation into biomass. Yet, ^13^C-labeling experiments conclusively showed that formate was not being assimilated. These results remain unexplained. One possible explanation could be that the pathway used by the *ΔserA* strains to synthesize serine has a higher energy demand, and the oxidation of formate may supply the necessary energy to support normal growth levels. Such increase in biomass formation due to increased energy availability have been reported in *P. putida* strains supplemented with formate^43^. However, further investigation would be needed to resolve this discrepancy.

While we did not achieve successful engineering of formate assimilation via the rGlyP in *P. thermoglucosidasius*, we believe the findings presented in this study offer valuable insights and serve as a foundation for future research in this area.

## Materials and methods

### General cultivation techniques, growth curves determination, auxotrophy investigation of the Δ*serA* strain, media composition, bacterial strains and plasmids

For the cultivation of *E. coli* strains, lysogeny broth (LB) medium was used containing 10 g/L tryptone (Bacto^TM^), 10 g/L NaCl, and 5 g/L yeast extract (Bacto^TM^). Solid LB medium additionally contained 15 g/L agar. *E. coli* strains were generally cultivated in 10 mL liquid medium in falcon tubes at 37°C and 200 rpm overnight or on solid medium at 37°C overnight.

For general cultivation of *P. thermoglucosidasius* strains, TGP medium was used containing 17 g/L tryptone (Bacto^TM^), 3 g/L peptone (Sigma Aldrich), 5 g/L NaCl, 2.5 g/L K_2_HPO_4_, 0.4% w/v glycerol, and 4 g/L sodium pyruvate. Solid TGP medium additionally contained 30 g/L agar.

For the experimental determination of growth curves of *P. thermoglucosidasius*, the auxotrophic phenotype investigation of the Δ*serA* strain, and the validation of formate assimilation, thermophilic minimal media (TMM) was used^44^. TMM contained: 494 mg/L NaCl, 145 mg/L NaSO_4_, 24.7 mg/L KCl, 3.97 mg/L KBr, 184 mg/L MgCl_2_·6 H2O, 89.2 mg/L NaNO_3_, 8.37 g/L MOPS, 3.58 mg/L FeSO_4_, 716 mg/L tricine, 229 mg/L K_2_HPO_4_, 0.51 g/L NH_4_Cl, 73.5 mg/L CaCl_2_, 2 μg/L biotin, 5 μg/L thiamine-HCl, 2 μg/L folic acid, 0.1ug/L vitamin B12, 0.127 μg/L spermidine, 174 μg/L spermine tetrahydrochloride, 0.7 mg/L hemin, 1 mg/L FeCl_3_·6 H2O, 0.18 mg/L ZnSO_4_·7 H2O, 0.12 mg/L CuCl_2_·2 H2O, 0.12 mg/L, MnSO_4_· H2O, and 0.18 mg/L CoCl_2_·6 H2O, with a variable carbon source.

For the determination of growth curves of *P. thermoglucosidasius* strains, cells were inoculated from glycerol stocks and precultured overnight in 50 mL falcon tubes containing 10 mL of TGP media, at 60°C and 180 rpm in a shaking incubator (New Brunswick™ Innova® 44 Series). After an overnight incubation, cell optical density of the bacterial precultures was measured with a portable Implen™ OD_600_ DiluPhotometer, the preculture media was removed by centrifugation and decantation and the cells were resuspended in TMM. These cell suspensions were used as inoculum for the determination of growth curves and the amount of cells harvested was corrected depending on the OD_600_ of the precultures, to yield an initial OD_600_ ~ 0.4 in the microtiter well. The strains were inoculated in quintuplicates and cultivated in microtiter flat-bottom 96-well plates, in TMM media with either glucose (5 g/L) or a mixture of glucose (5 g/L) and formate (1.35 g/L) as carbon source. Culture volume per well was 150 µL. To avoid evaporation, 50 µL of mineral oil (Sigma-Aldrich) were added on each well. These experiments were performed in a BioTek Synergy Neo2 Reader (Agilent) operated at 57°C, with 10% CO_2_ and 90% air, while alternating between one minute orbital shaking and one minute linear shaking (1 mm amplitude) and measuring OD_600_ every 12 minutes.

For the investigation of the auxotrophic phenotype of the Δ*serA* strain, precultures were prepared in TGP as explained in the previous paragraph. After an overnight incubation, cell optical density of the bacterial precultures was measured the OD_600_, the preculture media was removed by centrifugation and decantation and the cells were resuspended in 50 mL falcon tubes containing 10 mL of TMM medium supplemented with either serine (210 mg/L) or glycine (150 mg/L), along with glucose (2 g/L) as carbon source, at an initial OD_600_~0.05. Cultures were inoculated in triplicates and incubated at 60°C and 180 rpm, and OD_600_ was measured twice, 24 and 48 h after inoculation.

When needed for cloning or plasmid propagation, chloramphenicol (15 µg/L) was added to *E. coli* cultures, while thiamphenicol (6.5 µg/L) was added to *P. thermoglucosidasius* cultures.

### Plasmid assembly and genome modification

Generally, primers were designed to either introduce BsaI sites and overhangs for Golden Gate assembly or to provide homology for Gibson assembly (Table S1). All the desired DNA fragments were amplified using Q5 polymerase (Q5 2x MasterMix, New England Biolabs), isolated from a 1% agarose gel (Zymogen gel DNA recovery kit) and assembled by either Gibson (50 °C, 1 hour) or Golden Gate ((37°C, 5 min; 16 °C, 5 min) × 30; 37°C, 20 min; 65°C, 20 min). 5 µL of assembly mixture were transformed by heat shock in Neb5a commercial chemically competent cells (New England Biolabs) and plasmid presence in bacterial cells was verified by colony PCR using OneTaq polymerase (OneTaq 2x MM, New England Biolabs). Plasmids were isolated (QIAprep Spin Miniprep Kit, Qiagen) and the sequence of all newly assembled plasmids were confirmed by Oxford Nanopore whole plasmid sequencing (Plasmidsaurus).

To obtain the plasmid pJS15A, a derivative of the Geobacilli: *E. coli* shuttle vector pJS28 (Table 2) we replaced by Golden Gate assembly the origin ColE1 from pJS28 by a lower copy number p15A ori ^45^. To construct the plasmids pJS15A-*glyA*KO and pJS15A-*serA*KO, first *thermoCas9* under the expression of the *P. thermoglucosidasius* native cellobiose inducible promoter P*bgl*, and the sgRNA module including a scrambled spacer were amplified by PCR and assembled into the backbone pJS15A by Golden Gate. The resulting vector was later digested with SacI, and 1 KB homology arms (HA) flanking the upstream and downstream regions of either the *glyA* or *serA* gene of *P. thermoglucosidasius* were included via Gibson assembly. The scrambled spacer was replaced by a targeting spacer via Gibson assembly.

Confirmed plasmids were transformed into *P. thermoglucosidasius* by conjugation, following a slightly adapted protocol from Hon Wai Wan (2013)^46^. Notable differences were that a 50 mL culture of *P. thermoglucosidasius* was grown to mid-log phase (~0.7 OD_600_) in a 250 mL baffled flask and then centrifuged at 4500xg for 10 minutes before the actual conjugation.

To delete the genes *glyA* and *serA*, we induced homologous recombination of HA from pJS15A-*thermocas9*-*glyA*KO or pJS15A-*thermocas9*-*serA*KO onto the genome of *P. thermoglucosidasius* WT strain and selected against wild-type DNA by using ThermoCas9. One confirmed transformant of *P. thermoglucosidasius* containing the editing plasmid was cultivated in 10 mL liquid TGP containing chloramphenicol (8 µg/mL) at 60°C overnight, then transferred (1mL) to fresh 10 mL TGP containing chloramphenicol (8 µg/mL) and cultivated at 68°C overnight. The resulting culture was then transferred into fresh TGP containing chloramphenicol (8 µg/mL) and 1% w/v cellobiose and cultivated at 60°C overnight to induce the P*Bgl* promoter and enhance *thermocas9* transcription. Finally, the culture was plated on a TGP plate with 1% w/v cellobiose using 1 mL, 100 µL, 10 µL, or 1 µL of liquid culture and cultivated at 55°C. Gene deletions were confirmed by colony PCR.

### Validation of formate assimilation via ^13^C labelling experiments

To validate formate assimilation into biomass, we performed steady-state labelling experiments with ^13^C labelled formate. *P. thermoglucosidasius* WT-C1MT and Δ*serA*-C1MT strains were inoculated from glycerol stocks in liquid TMM media supplemented with 5 g/L of glucose, 1.38 g/L of ^13^C labelled formate. The precultures were incubated overnight at 52°C, 180 rpm, 10% CO_2_ and 90% air. 100 µL from each preculture were subcultured in 10 mL of fresh media. Cells were incubated at 52°C and 180 rpm, 10% CO_2_, grown to late exponential phase (OD~1) and one mL of culture broth was harvested and centrifuged (13000 rpm, 1 minute). All cultivations were performed in a CO_2_ incubator S41i (Eppendorf). Cell pellets were stored at −20 until further processing. Pellets were re-suspended in 200 µL of 6 M HCl and incubated at 110 °C for 16 hours to lyse the cells and hydrolyze the respective proteomes. Hydrolyzed samples were then filtered on a filter plate (AcroPrep Advance 96-well Filter, 0.2 µm membrane, product ID 8019) by positive pressure, and dried in a cold-trap speedvac. Dried samples were then derivatized by adding 35 µL of pyridine (Sigma-Aldrich, cat no. 270407) and 50 µL of MBTSFA + 1% (wt/wt) TBDMSClS (Sigma-Aldrich, cat no. 375934), mixed by pipetting and incubated at 60 °C for 30 min. 75 µL of the derivatized samples were aliquoted to LC-MS vials with glass inserts for GC-MS analysis within 12h from derivatization on an Agilent 5977 GC-MS system with an Agilent DB-5ms capillary column (30m, inner diameter of 0.25 mm, film thickness of 0.25 µm, cat no. 122-5532). Samples were measured in full-scan mode, using a 1:100 split ratio, with the following gradient: start at 160 °C, hold for 2 min, ramp to 310 °C at 7 °C/min, hold for 1 min. We considered for further analysis fragments listed in Long C.P. & Antoniewicz (2019) which included the whole C backbone of the relative amino acid (Gly 246, Ser 390)^49^. Raw chromatographic data was integrated using SmartPeak^50^. The relative data was corrected for the natural abundance of isotopes in the derivatization agents used for GCMS analysis^51^.

