## Supplementary data for "*Parageobacillus thermoglucosidasius* and the reductive glycine pathway: A journey of Almosts and Maybes"

### Chapter 5

#### Supplementary materials

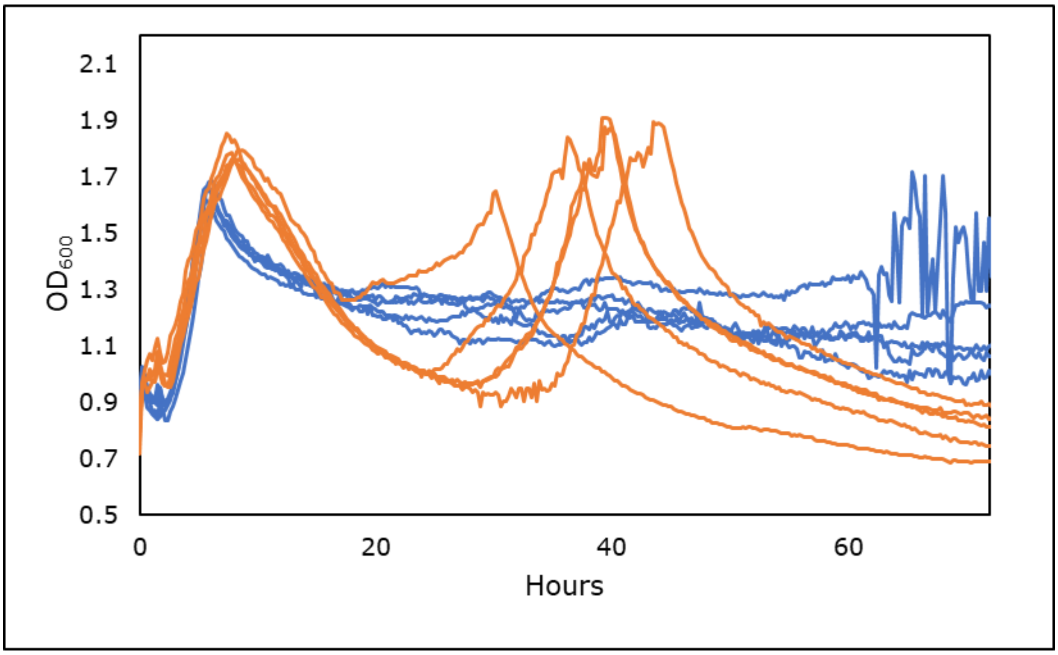


Figure S1: Growth curves of *P. thermoglucosidasius* WT-EV, when either glucose (5 g/L) or a mixture of glucose (5 g/L) and formate (1.35 g/L) were used as carbon source. Individual curves represent one replicate. Orange lines = Glucose. Blue lines= Glucose and formate.

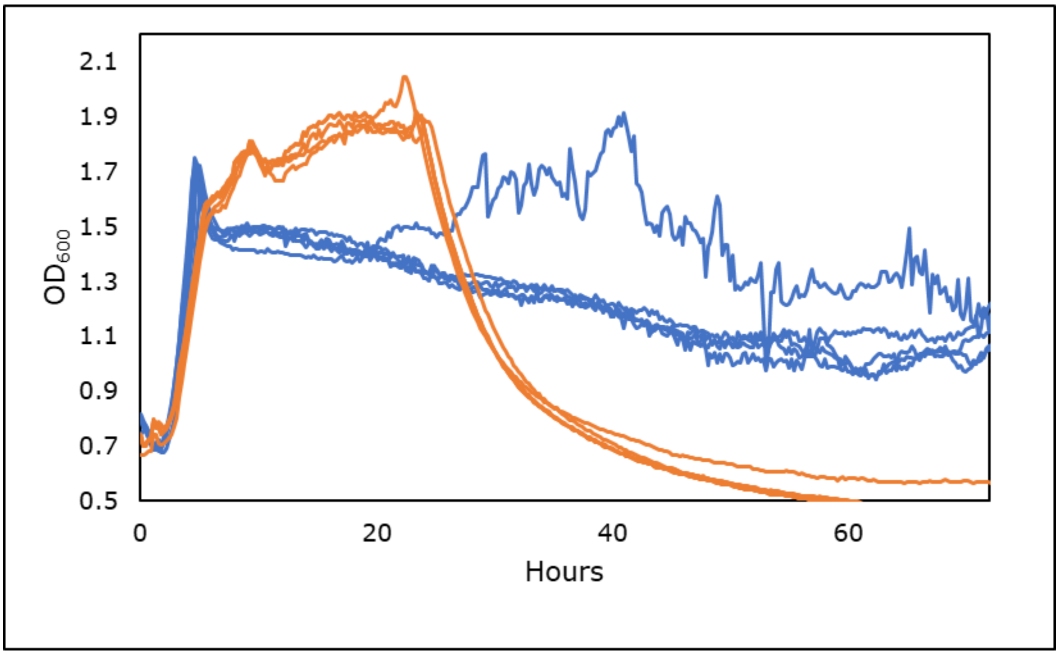


Figure S2: Growth curves of *P. thermoglucosidasius* WT-C1MT, when either glucose (5 g/L) or a mixture of glucose (5 g/L) and formate (1.35 g/L) were used as carbon source. Individual curves represent one replicate. Orange lines = Glucose. Blue lines= Glucose and formate.

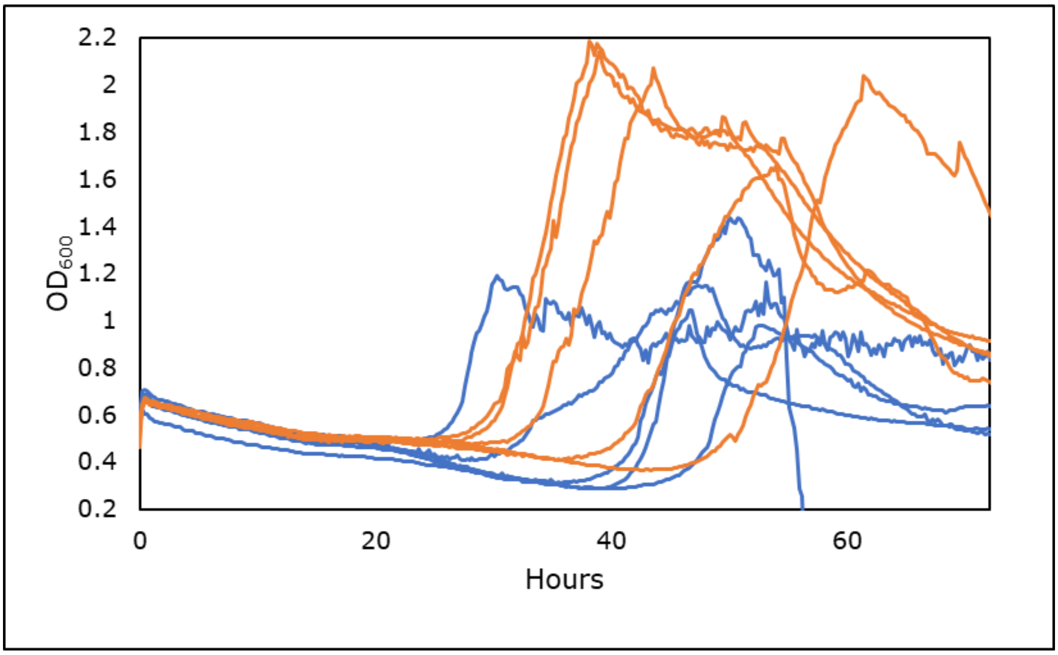


Figure S3: Growth curves of *P. thermoglucosidasius ΔserA*-EV, when either glucose (5 g/L) or a mixture of glucose (5 g/L) and formate (1.35 g/L) were used as carbon source. Individual curves represent one replicate. Orange lines = Glucose. Blue lines= Glucose and formate.

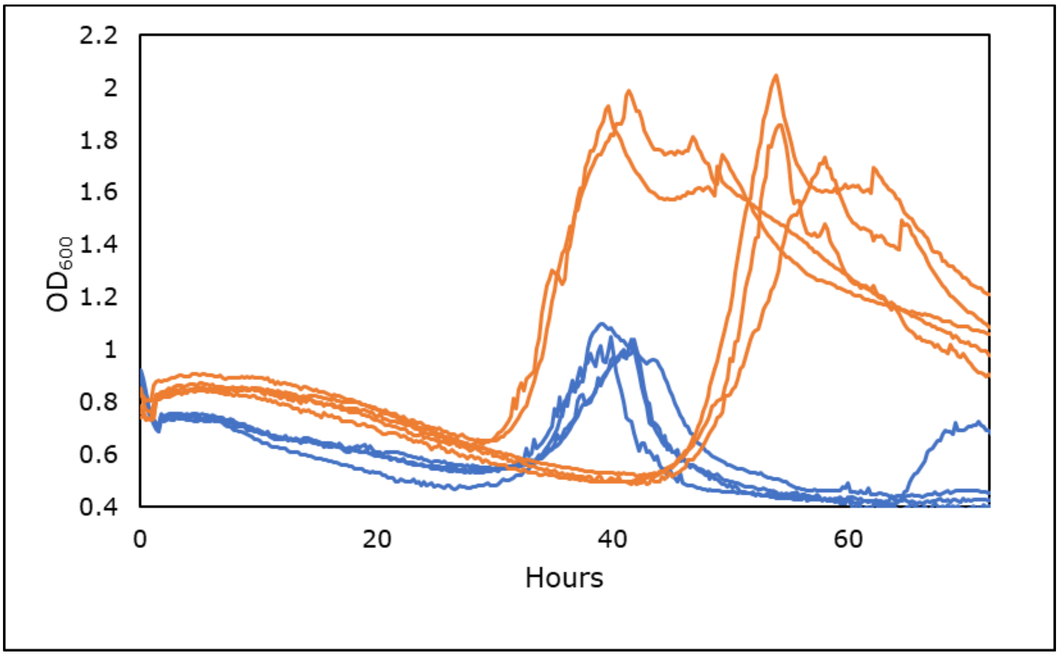


Figure S4: Growth curves of *P. thermoglucosidasius ΔserA*-C1MT, when either glucose (5 g/L) or a mixture of glucose (5 g/L) and formate (1.35 g/L) were used as carbon source. Individual curves represent one replicate. Orange lines = Glucose. Blue lines= Glucose and formate.

| Name | Sequence |
|---|---|
| pJS28-part1-GG-F | tggtctcaatggtcatagctgtttcctgtgtg |
| pJS28-part1-GG-R | tggtctcatagaccatggagagaaaagaaaatcg |
| pJS28-part2-GG-F | tggtctcatctaacttttccactttttgtcttgtcc |
| pJS28-part2-GG-R | tggtctcagcgcggtatcattgcagc |
| pJS28-part3-GG-F | tggtctcagcgcgacccacgctcaccgg |
| pJS28-part3-GG-R | tggtctcagcatgcctgcaggtcgac |
| pJS15A_BsaI_F | tggtctcatcatagctgtttcctgtgtgaa |
| pJS15A_BsaI_R | tggtctcagcatgcctgcaggtc |
| Pbgl_GG_F | tggtctcaatgccgataaacgcgaagaaggtg |
| Pbgl_GG_R | tggtctcatcgaatcactccttatctagaatgcaacctcctttatgttcg |
| TCas9_part1_GG_F | tggtctcctcgaatgaagtataaaatcggtcttgatatc |
| TCas9_part1_GG_R | aggtctcatggtgagccgtactttttg |
| TCas9_part2_GG_F | aggtctcaaccagtctccatccatatcgaactg |
| TCas9_part2_GG_R | tggtctcaccatgattacgccaagcttc |
| USHR_glyASacI_F | CTGATTGTGAAATTGAATTCGAGCTatcccgattacgcgattcc |
| USHR_glyAKO_R | ccccctcgtttcgaaattaagaatattc |
| DSHR_glyAKO_F | tattcttaatttcgaaacgagggggcgaacacgcgcctctaaag |
| DSHR_glyASacI_R | GGCAAGGTCATGATGGGCGTGGTCGGGTACCGagctttcgcttgaaaaccttatcacatatcc |
| USHR_serASacI_F | CTGATTGTGAAATTGAATTCGAGCTatcaaagtcgcagacgagc |
| USHR_serAKO_R | aaccagaagccttttaatgaaatggatggtgagctcctttctttaacaag |
| DSHR_serAKO_F | agaaaggagctcaccatccatttcattaaaaaggcttctggttacc |
| DSHR_serASacI_R | GATGGGCGTGGTCGGGTACCGagctaggttttaaagatattaaaaatctggataaggaag |
| USHR_aceAKO_F | CTGATTGTGAAATTGAATTCGAGCTgcaaaaaaatggcgccg |
| USHR_aceAKO_R | ccccttctcttttatataaactccccatatccccttttccttccccatccttattg |
| DSHR_aceAKO_F | aggggatatgggggagtttatataaaag |
| DSHR_aceAKO_R | GATGGGCGTGGTCGGGTACCGagcttgaaggtgagaaatccatacgc |
| Protospacer_glyA_R | gagcatgcgaacgtacagccgcaGTATAACGGTATCCATTTTAAGAATAATCCATTTTTC |
| Protospacer_glyA_F | ACCGTTATACtgcggctgtacgttcgcatgctcGTCATAGTTCCCCTGAGATTATCG |
| Protospacer_serA_R | ttagaaaggagttcttcggtcacGTATAACGGTATCCATTTTAAGAATAATCCATTTTTC |
| Protospacer_serA_F | ACCGTTATACgtgaccgaagaactcctttctaaGTCATAGTTCCCCTGAGATTATCG |
| Protospacer_aceA_R | gatacatttgcccggcaaggttaGTATAACGGTATCCATTTTAAGAATAATCCATTTTTC |
| Protospacer_aceA_F | ACCGTTATACtaaccttgccgggcaaatgtatcGTCATAGTTCCCCTGAGATTATCGC |
| glyA-KO-check-F | gaaagggtaacgttggtttttcc |
| glyA-KO-check-R | GTCTGATCACGCAGGAATTCATAT |
| serA-KO-check-F | cctgtggtgcatgtgg |
| serA-KO-check-R | TGGGGGATGCGTTATGAG |
| aceA-KO-check-F | CGTCCACATTTGGACGTTTTTTAC |
| aceA-KO-check-R | TAGACGATCGTCTGGTCATTTTTTG |
| pLDHgst_gen_F | CTGATTGTGAAATTGAATTCGAGCTGCGGGACGGGAGC |
| pLDHgst_gen_R | CATTCGAATCACTCCTTATCTAGAATCCTCC |

| | |
|---|---|
| fhsMT_pLDH_F | GGAGGATTCTAGATAAGGAGTGATTCGAATGttgtccaaggtacccagtg |
| fhsMT_R | ctgggctggcatTCGAATCACTCCTTATCTAGActagaaaagaccggtaatgacgc |
| folDMT_F | cgtcattaccggtcttttctagTCTAGATAAGGAGTGATTCGAatgccagcccagatcc |
| folDMT_R | GATGGGCGTGGTCGGGTACCGagctctagcgccgggcc |

Table S1: List of all primers used.
